# Resolving Taxonomic Boundaries and Revealing Genetic Diversity in *Enterococcus casseliflavus* and Closely Related Taxa

**DOI:** 10.1101/2024.09.16.613146

**Authors:** Matheus M. S. M. Lima, Janira Prichula, Tetsu Sakamoto

## Abstract

*Enterococcus casseliflavus* is a commensal bacterium that can occasionally cause human infections. A concern is that all strains of this species carry the *vanC* operon on their chromosomes, which reduces susceptibility to vancomycin. Aside from that, the classification of *E. casseliflavus* remains challenging because it shares up to 99% 16S rRNA sequence identity with other enterococci species. Here, we reassess the taxonomy and genomic diversity of *E. casseliflavus* and closely related taxa by comprehensively analyzing publicly available genomes to clarify their evolutionary relationships, ecological distributions, and clinical relevance. We retrieved 156 genomes of clinical and non-clinical isolates that showed ANI >90% with *E. casseliflavus* from public databases. Using ANI, core genome-based phylogeny, and pangenome reconstruction, we consistently identified three well-defined clusters corresponding to *E. casseliflavus*, *E. entomosocium*, and *E. innesii*. Our results indicate that a higher ANI threshold (>96.6%) is required to accurately discriminate between species within this group, as many strains showed more than 95% ANI with multiple species. Our results also support that *E. flavescens* and *E. casseliflavus* represent a single species and should be treated as synonyms. Genes related to biocide resistance, metal tolerance, and virulence were identified across the three species. *E. casseliflavus* exhibited the greatest diversity of resistance determinants. All species harbored the *vanC* operon associated with intrinsic vancomycin resistance, while two genomes additionally carried the *vanD* operon linked to high-level glycopeptide resistance. Overall, our findings refine species boundaries and highlight the importance of genome-based approaches for accurate classification and surveillance within the *Enterococcus* genus.

## Introduction

*Enterococcus* is a Gram-positive bacterium that emerged around 400 – 500 million years ago, coinciding with the emergence of animal territorialization, and has become a common component of the gut microbiomes of land animals (Lebreton et al. 2017). While primarily found in the guts of humans and other animals (Huff et al. 2026; Lebreton et al., 2014; Mocellin et al. 2024; Prichula et al. 2021; Schwartzman et al. 2024), in the past three decades, enterococci have also become an opportunistic pathogen, one of the major causes of multidrug-resistant hospital-associated infections (Lebreton et al. 2017; Miller and Arias 2024), including urinary tract infections, endocarditis, and bacteremia, primarily due to vancomycin-resistant *E. faecalis* and *E. faecium* (*Centers for Disease Control and Prevention (CDC); European Centre for Disease Prevention and Control - ECDC (2024)*; Ikuta et al. 2022; Lebreton et al. 2018; Miller and Arias 2024). Currently, more than 90 species of *Enterococcus* are described, with a valid published name, including synonyms [(Parte et al. 2020); LPSN, consulted on 04/10/2026)]. The phylogenetic studies of the genus divided them into four clades: I, II, III, and IV (Lebreton et al. 2017; Schwartzman et al. 2024).

*Enterococcus casseliflavus* belongs to Clade IV, is a mobile, yellow-colored bacterium that occurs naturally in the guts of land and marine animals (Lebreton et al., 2017; Prichula et al., 2021; Schwartzman et al. 2024). However, *E. casseliflavus* has been occasionally isolated from human cases of bacteremia and other infections, which are of particular concern, as all strains described so far harbor the *VanC* gene cluster, which reduces susceptibility to vancomycin, one of the few antibiotics that are bactericidal for enterococci (Collins et al., 1984; Palmer et al., 2012; Pappas et al., 2004; Prichula et al., 2021; Schwartzman et al., 2024; Vasilakopoulou et al., 2020). As an opportunistic pathogen in hospital infections (Phute, et al. 2025; Yoshino 2023), it is associated with endocarditis (Okumura et al. 2021) and bacteremia (Fujii et al. 2024; Mohammed and Yeung 2024; Vasilakopoulou et al. 2020), endophthalmitis (Kandarakis et al. 2023; Priyadarshini et al. 2025), and bacterial peritonitis (Yin et al. 2018). Furthermore, there are reports of this species in patients with serious comorbidities, such as hematologic cancer, organ transplant recipients, kidney failure, diabetes mellitus, and vascular and intravenous infections (Phute, et al. 2025; Vasilakopoulou et al. 2020; Xu et al. 2024).

Since the description of *E. casseliflavus* in 1984 (Vaughan et al. 1979; Collins et al., 1984), other species close to it have been described. In 1992, *E. flavescens* was described as a new species closely related to *E. casseliflavus*, distinguished by its inability to ferment ribose and by the absence of hemolysis on sheep blood (Pompei et al. 1992). However, subsequent molecular studies have reported difficulties in distinguishing the two species. Descheemaeker and colleagues (Descheemaeker et al. 1997) found that it was not possible to differentiate the two species using pulsed-field gel electrophoresis (PFGE) and PCR-based typing. Studies comparing *16S rRNA* gene sequences have demonstrated 100% identity between the two species (Patel et al., 1998). Furthermore, DNA-DNA hybridization analyses also suggested that *E. casseliflavus* and *E. flavescens* were the same species (Teixeira et al. 1997). Other studies have also compared genetic markers between the two species and consistently found high levels of similarity (Baele et al. 2000; Dutka-Malen et al. 1995; Navarro and Courvalin 1994; Poyart et al. 2000; Quednau et al. 1998; Svec et al. 2005). Naser and colleagues (Naser et al. 2006) showed the close relationship between the two species by comparing three core genes of the two species, phenylalanyl-tRNA synthetase alpha subunit (*pheS*), RNA polymerase alpha subunit (*rpoA*) and the alpha subunit of ATP synthase (*atpA*), as well as performing DNA-DNA hybridization experiments. Therefore, they proposed that, under the principle of priority, *E. casseliflavus* should be retained and *E. flavescens* should be reclassified as a synonym (Naser et al. 2006). This classification is still maintained according to the List of Prokaryotic names with Standing in Nomenclature – LPSN (Parte et al. 2020).

Recently, two new species have been described as genetically close to *E. casseliflavus*. One such species, *E. innesii* (Gooch et al. 2021), was isolated from the wax moth *Galleria mellonella*. It exhibits digital DNA–DNA hybridization (dDDH) values of 59% and an average nucleotide identity (ANI) of 94.5% relative to *E. casseliflavus* NBRC 100478, its closest relative, supporting its classification as a distinct species based on established genomic thresholds (≥95%; Jain et al. 2018). A second species, *E. entomosocium* (Gomes et al. 2023), was isolated from the intestinal tract of *Spodoptera frugiperda* larvae and is notable for its potential role in pesticide degradation. Its designation as a novel species was supported by a combination of phenotypic characterization and comparative genomics analyses, revealing ANI values of 94.7% relative to *E. casseliflavus* EC20 (Gomes et al. 2023). To date, no comprehensive study has systematically revised the taxonomic classification and the genetic diversity of *E. casseliflavus* and its closely related species. Here, we address this gap by analyzing genomic data of 156 strains from public domains, including both clinical and non-clinical strains of *E. casseliflavus* and closely related taxa. Through a comprehensive analysis of publicly available genomes, we clarify their evolutionary relationships, ecological distribution, and clinical relevance.

## Material and Methods

### Genome collection and data processing workflow

*Enterococcus* genomes available in NCBI’s RefSeq database (*National Center for Biotechnology Information - NCBI*) were used for this study. A total of 8,976 genomes belonging to the genus were accessed on June 19, 2024. First, the Average Nucleotide Identity (ANI) of these genomes was analyzed using FastANI (Jain et al. 2018). Genomes with 90% or higher compared to a type strain of *Enterococcus casseliflavus* DSM 20680 (ATCC 25788; reference genome GA: GCF_001885845.1) (Collins et al. 1984; Vaughan et al. 1979), were selected. Genomes were further filtered to *i)* remove low-quality assemblies, *ii)* redundant sequences from the same strain, and *iii)* ensure representative sampling across sources. Finally, the genomes were annotated using PROKKA (Seemann 2014) with default parameters to ensure uniform annotation.

### All-to-all pairwise Average Nucleotide Identity (ANI)

To understand the relationships among the selected genomes, an all-to-all pairwise ANI analysis was performed using FastANI. Type-strains *Enterococcu*s *innesii* GAL7 [GA: GCF_018982785.1; (Gooch et al. 2021)], *E. entomosocium* IIL-Cl05 [GA: GCF_030441895.1; (Gomes et al. 2023), and *E. casseliflavus* DSM 20680 [ATCC 25788, GA: GCF_001885845.1; (Collins et al. 1984; Vaughan et al. 1979)] were considered as representative genomes of each species. Additionally, *E. flavescens* ATCC 49996 [GA: GCF_000407405.1; (Pompei et al. 1992)] strain was included as a representative of *E. flavescens*, which is considered a synonymous species of *E. casseliflavus* (Parte et al. 2020). To visually represent the nucleotide identity between these genomes, a heatmap was generated using the packages reshape, ggplots, and ComplexHeatmap (Gu, 2022) from R.

### Analysis of Genomic features, resistome and virulome

Putative determinants of resistance to biocides and metal tolerance, virulence, and antimicrobial resistance genes were predicted using PanViTa (Rodrigues et al., 2023). This tool uses the BacMet2 (antibacterial biocide and metal resistance genes database; Pal et al., 2014), the VFDB (Virulence Factor Database; Liu et al., 2022), and the CARD (Comprehensive Antibiotic Resistance Database; Alcock et al., 2023) databases to perform the predictions. All predictions were performed with standard thresholds of 70% identity and 70% coverage.

### Pangenome analysis and core genome-based phylogeny

The pangenome of all samples was determined using the ortholog groups identified by Orthofinder (Emms and Kelly 2019), employing files annotated by PROKKA (Seemann 2014) as input. The multiple alignment of the core genes generated by Orthofinder was used to perform a phylogenomic analysis. This was performed by FastTree software (Price et al. 2010) using the GT+C substitution model, and 1000 bootstrap samples. The resulting phylogenetic tree was rooted at the midpoint, annotated using TaxOnTree (Sakamoto and Ortega 2020), visualized with FigTree (http://tree.bio.ed.ac.uk/software/figtree/), and edited with Inkscape (https://inkscape.org).

The rarefaction curves for pan gene accumulation and core gene reduction were generated using presence-absence data from ortholog groups identified by Orthofinder (Emms and Kelly 2019), combined with an in-house Python script. Genomes in the dataset were sampled sequentially, and the numbers of pan and core genes were recorded as each genome was added. This process was repeated 1,000 times. The mean number of pan and core genes was then calculated to fit power-law regressions based on Heap’s law (Tettelin et al., 2008), estimating the growth (γ) and reduction (σ) rates of pan and core genes, respectively. Model fitness was assessed using the coefficient of determination (R^2^), calculated by first determining the R^2^ values between the model’s predicted data and the data of each iteration of the rarefaction analysis, then computing the mean R^2^.

### Statistical analysis

We used non-parametric statistical methods to evaluate differences in genomic features (number of genes, genome size, and GC content) among groups defined by species and isolation source. The Kruskal–Wallis H-test (Kruskal and Wallis, 1952) assessed overall differences in median values across groups. When significant differences were observed (p < 0.05), post hoc pairwise comparisons were conducted using Dunn’s test with p-values adjusted using the Holm correction method (Dunn, 1961; Holm, 1979) to identify specific group differences. All statistical analyses were performed in Python. The Kruskal–Wallis test was performed using the Kruskal function from the SciPy library (Virtanen et al. 2020), and Dunn’s test was carried out using the posthoc_dunn() function from the scikit-posthocs package (Terpilowski 2019).

## Results and Discussion

### Phylogenomic analysis reveals that a higher ANI threshold is required to discriminate E. casseliflavus and closely related species

A total of 8,976 reference genomes belonging to the genus were accessed on June 19, 2024. Among these, only 182 genomes exhibited 90% ANI or higher compared to *E. casseliflavus* type strain DSM 20680 [ATCC 25788, GA: GCF_001885845.1; (Collins et al. 1984)]. After removing low-quality assemblies, redundant sequences from the same strain, and ensuring representative sampling across sources, this set was further refined to 156 genomes (Table S1). According to the taxonomic annotation obtained from the NCBI database, of the 156 samples, 25 had only genus-level identification, while the others belonged to the species *E. casseliflavus* (n=115), *E. entomosocium* (n=3), and *E. innesii* (n=13). The sizes of the analyzed genomes ranged from 3.2 to 4.1 Mb, with a GC content between 40.2% and 43.3%. The number of sequence components (contigs and scaffolds) varied from 1 to 199, and sequencing coverage ranged from 22.2x to 2,571x. The genomes included in this study were isolated between 1984 and 2023 (Table S1).

To verify the relationship across the 156 selected genomes, an all-to-all pairwise ANI analysis was performed using FastANI (Table S2; Supplementary Fig. 1). This analysis revealed three distinct clusters (1–3) that diverged from the NCBI genus-level identification (Supplementary Fig. 1). The largest cluster (Cluster 1) comprised 112 strains, including the DSM 20680 strain, the type strain for *E. casseliflavus*, and the type strain for *E. flavescens* (ATCC 49996). Cluster 2 showed 30 strains grouped with *E. innesii* type strain (GAL7). The smallest cluster (Cluster 3) contained 14 strains, including the type strain *E. entomosocium* IIL-Cl05.

Average nucleotide identity (ANI) analyses indicate a clear genomic separation between *Enterococcus* and related genera, such as *Vagococcus* and *Carnobacterium* (<63% ANI), while distinct *Enterococcus* species usually differ by more than 5% ANI from their closest sister taxa (Lebreton et al. 2017). To delineate enterococci species, we analyzed the ANI values of the 156 genomes against these four type strains: *E. casseliflavus* DSM 20680, *E. flavescens* ATCC 49996, *E. innesii* GAL7, and *E. entomosocium* IIL-Cl05 (Table S3). The standard ANI cutoff (≥95%; Jain et al., 2018) was initially used; however, this threshold proved inadequate, as some strains exhibited ≥95% ANI to multiple reference species, preventing reliable discrimination of these three *Enterococcus* species (Table 1; Table S3).

**Table 1:** Number of genomes (n = 156) stratified by ANI values relative to each reference genome.

| Strain Type Genome | ANI<br>< 95 | ANI<br>95-96 | ANI<br>> 96 | ANI expect<br>$\geq 95$ |
| --- | --- | --- | --- | --- |
| *E. casseliflavus* DSM 20680 | 22 | 20 | 114 | 134 |
| *E. entomosocium* IIL-Cl05 | 76 | 65 | 15 | 80 |
| *E. flavescens* ATCC 49996 | 23 | 20 | 113 | 133 |
| *E. innesii* GAL7 | 106 | 20 | 30 | 50 |

Furthermore, an extreme case was observed in 21 strains that showed ANI values above 95% for all reference strains (Table S4), indicating that the standard 95% cutoff did not resolve these enterococci species (Table 1, Table S2). In this sense, we verified that a reasonable ANI value delineating the three *Enterococcus* species could be around 96.6% (Figure 1), since the lowest ANI value between a reference sample and a sample within the same cluster was 96.93%, while the highest ANI value between a reference sample and a sample from a different cluster was 96.27% (Table S2). An interesting question that warrants further investigation is the substantial variation in ANI values observed when a reference genome from one species is compared against genomes from other species. For instance, comparisons between *E. casseliflavus* DSM 20680 and *E. innesii* strains yielded ANI values ranging from 94.74% to 96.19%. Although a more stringent ANI threshold was required to distinguish the three species, a similar situation has been reported for *E. faecium* and *E. lactis*, for which a cutoff of 96% ANI was proposed to differentiate the two species (Daza et al. 2021).

**Figure 1:**
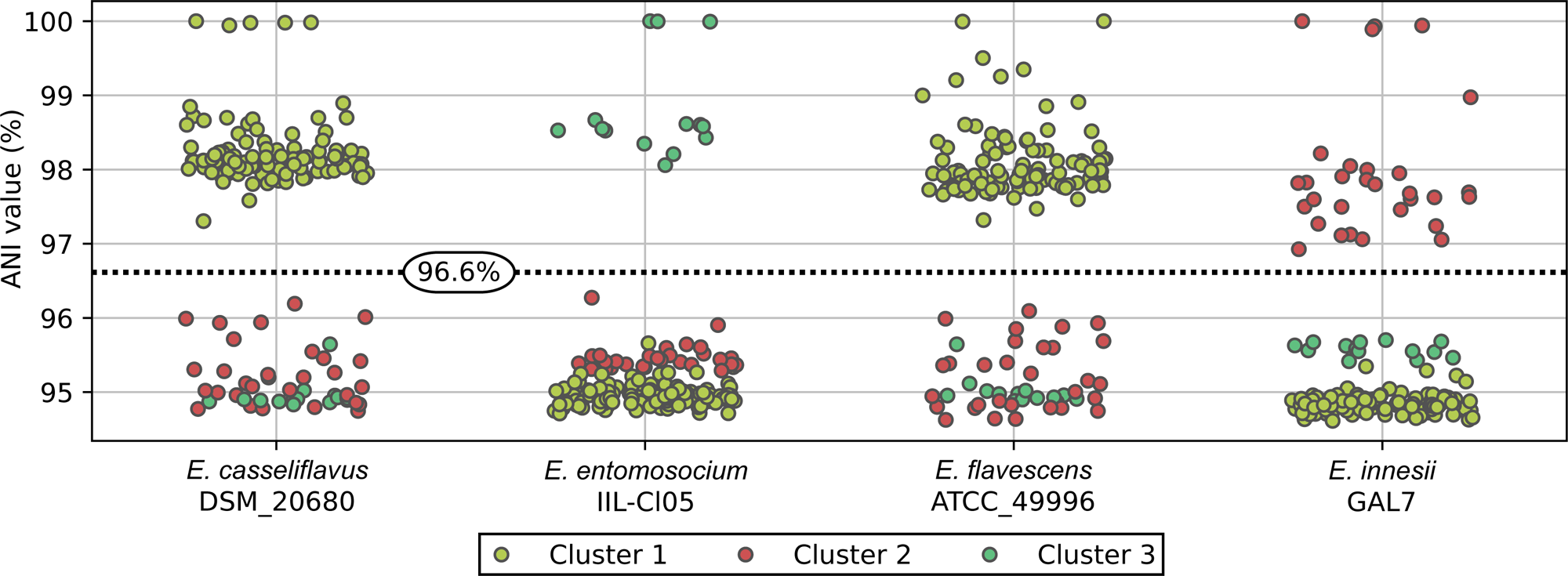
ANI values of 156 genomes relative to type strains. Colors indicate the cluster to which a sample belongs in the hierarchical clustering analysis using ANI values (Supplementary Fig. 1; Table S3). The dashed line indicates the suggested threshold (96.6%) for delineating the three *Enterococcus* species.

These results provide two key taxonomic insights: Cluster 1 (n=112), which includes the reference strains for *E. casseliflavus* and *E. flavescens*, supports their synonymy as a single species as suggested by Nasser and colleagues (Nasser et al., 2006). In contrast, Clusters 2 (n=30) and 3 (n=14) form distinct and well-resolved groups corresponding to *E. innesii* and *E. entomosocium*, respectively.

To further validate these findings, ortholog clustering was then conducted to identify single-copy core genes. To ensure consistency, all genomes were re-annotated using a standardized gene annotation pipeline (Seemann 2014). The clustering analysis of 156 genomes resulted in a total of 10,226 orthogroups (Table S5), of which 19.22% (1,966/10,226 orthogroups) were core genes, which represented, on average, 57.75% (1,966/3,404 genes) of the total genes of a sample (min: 48.26%; max: 67.37%). The phylogenomic tree inferred from single-copy core genes largely corroborated the ANI-based groupings (Figure 2), supporting the presence of the same three clusters identified in the previous analysis and reinforcing the need for taxonomic revision of the group based on an ANI threshold of >96.6%. These results highlight the importance of revisiting the reference genomes available in NCBI. Particularly, the strain ATCC 12755, which has been widely used as representative of *E. casseliflavus*, demonstrated to be consistently classified as *E. entomosocium* in this study.

**Figure 2:**
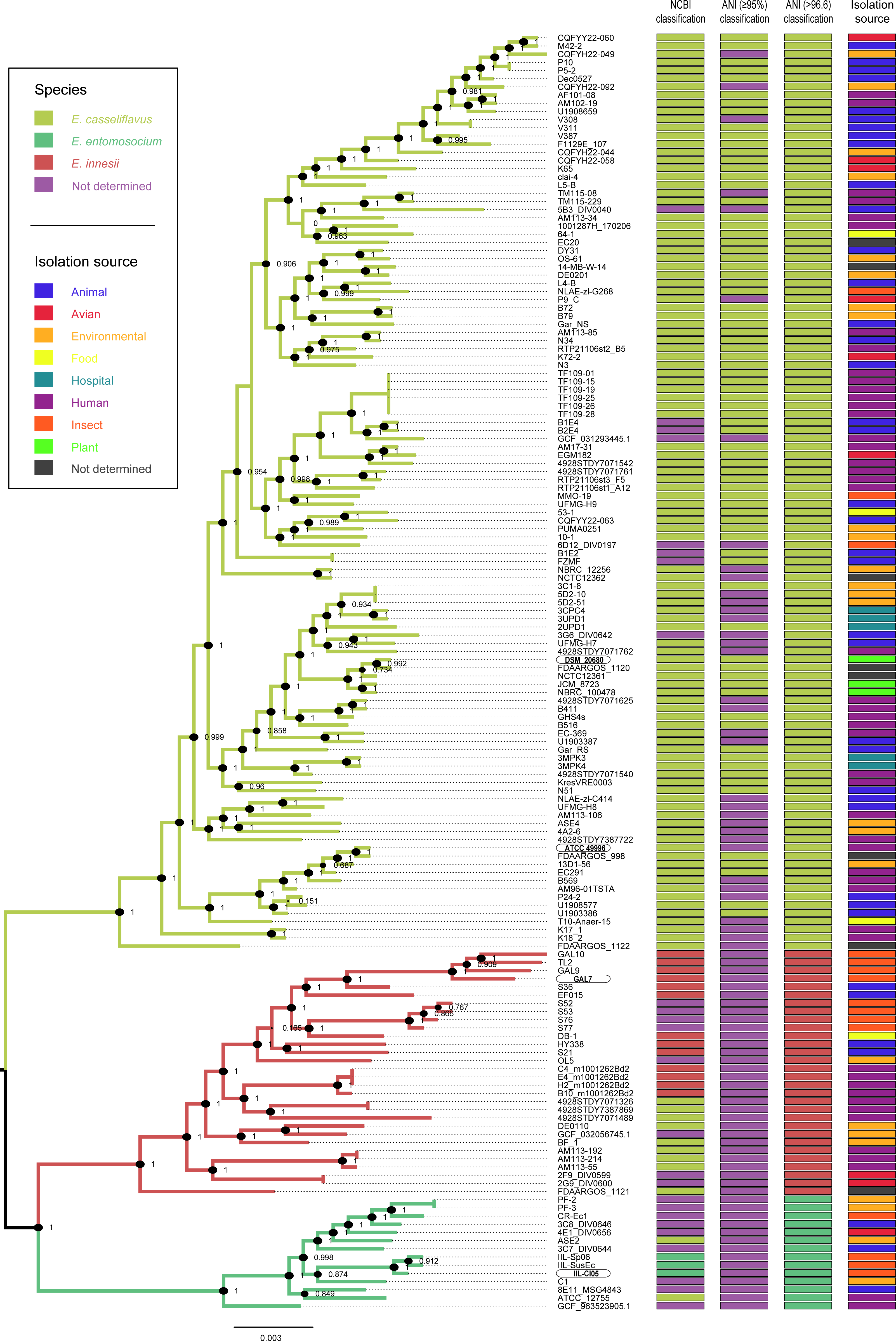
Phylogenomic tree of 156 *Enterococcus* spp. samples based on the 1,450 single-core genes. The three branches, identified by the colors, outline the species found among the samples. Species are highlighted in red (*E. innesii*), yellow (*E. casseliflavus*), and green (*E. entomosocium*). Four colored columns adjacent to the tree indicate, respectively, the species annotation based on NCBI classification, the ANI-based assignment for each sample [≥95%; (Jain et al. 2018); Table S3], the new classification proposed in this study (based on ANI >96.6% and core-genome analysis), and the source of isolation. Genomes that do not have species annotation in NCBI database, or that could not be assigned to a defined species (because strains exhibited ≥95% ANI to multiple reference species) are shown in purple. More details on the orthologous groups are provided in Table S5.

### Insights into the genomic features and pangenome of these three Enterococcus species

From this point, we understand the 156 genomes were partitioned into three groups using an ANI threshold of >96.6%, each corresponding to a distinct *Enterococcus* species: **(I)** *E. casseliflavus* (n = 112), comprising genomes with the highest identity to the DSM 20680 type strain; **(II)** *E. entomosocium* (n = 14) closely related genomes to IIL-Cl05 type strain; and **(III)** *E. innesii* (n = 30), consisting of genomes most closely related to the GAL7 type strain. We then characterized the genomic features and pangenome dynamics of these three species.

Across the *Enterococcus* genus, genome sizes range from ∼2.3 Mb in *E. sulfureus* (clade IV) to ∼5.4 Mb in *E. pallens* (clade III), with clade IV species averaging ∼3.0 Mb (Lebreton et al., 2017; Schwartzman et al., 2024). Consistent with this pattern, species in our dataset (Figure 3A) showed similar trends: *E. entomosocium* exhibited the largest genomes (mean 3.78 Mb; 3.26–4.14 Mb), followed by *E. innesii* (3.62 Mb; 3.26–3.93 Mb) and *E. casseliflavus* (3.58 Mb; 3.09–4.05 Mb). The Kruskal-Wallis test evidenced the existence of at least one pair of groups with different median genome size (p-value: 0.0011). The post-hoc Dunn test indicates significant differences between the genome size median of *E. casseliflavus* and *E. entomosocium* (adjusted p-value: 0.0010). Genome size among these species likely reflects differences in horizontal gene transfer (HGT), including gene gain and loss during evolution (Lebreton et al., 2017; Schwartzman et al., 2024; Zhong et al., 2017).

**Figure 3:**
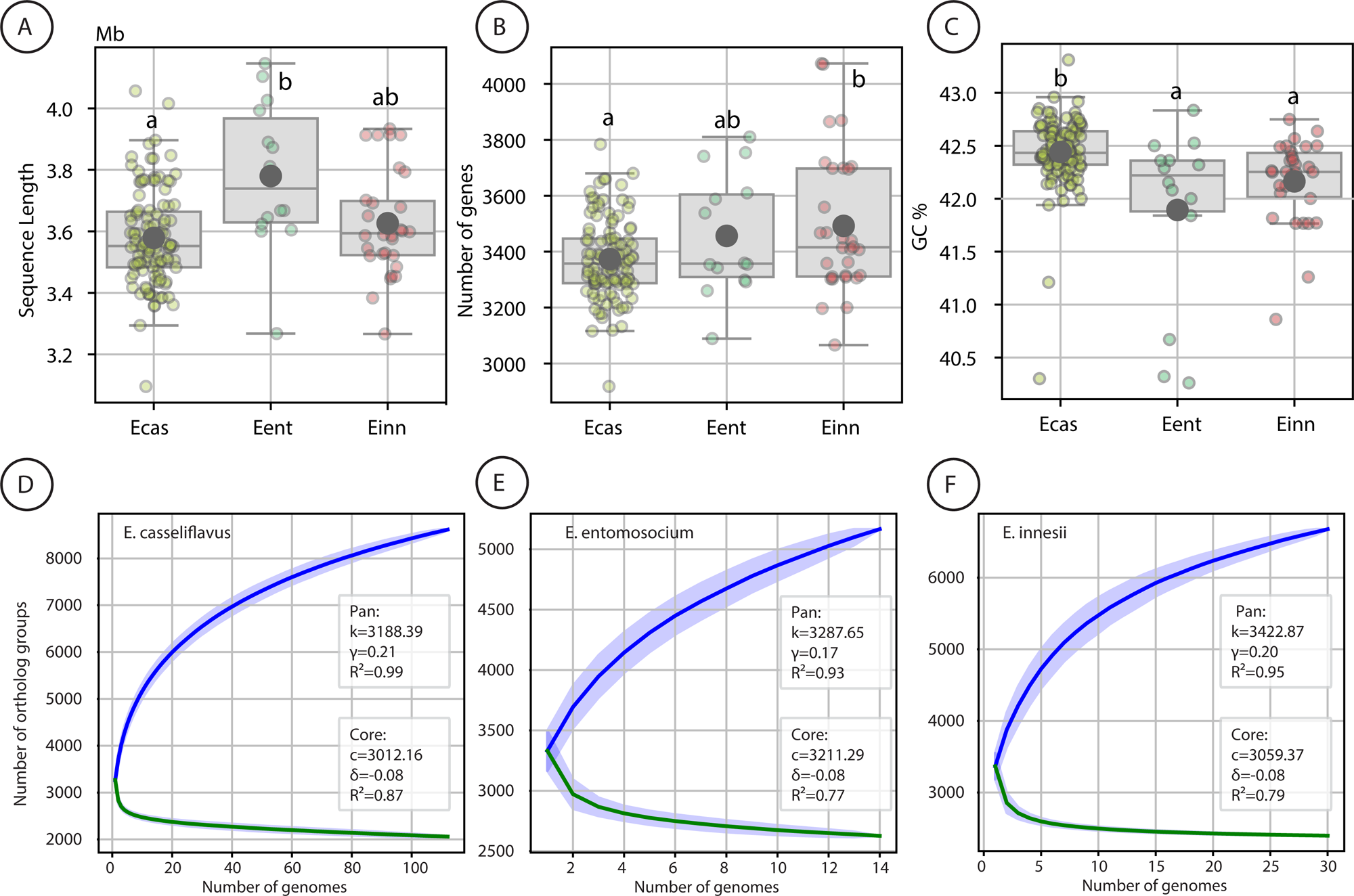
Genomic features and pangenome dynamics of three *Enterococcus* species. Boxplots comparing **(A)** genome size (in megabases), **(B)** the number of genes, and **(C)** GC content (%) among *E. casseliflavus* (Ecas), *E. entomosocium* (Eent), and *E. innesii* (Einn). Large gray circles indicate mean values. Different letters above the boxes denote statistically significant groupings based on Dunn’s test with Holm correction (q-value < 0.05). Pangenome and core genome curves for **(D)** *E. casseliflavus*, **(E)** *E. entomosocium*, and **(F)** *E. innesii*. Blue lines represent pangenome accumulation curves; green lines represent core genome reduction curves. Insets show the fitted models and parameters for each curve (k, γ for pangenome; C, δ for core genome), along with R² values indicating the goodness of fit.

Gene content varied among the three species (Figure 3B). *E. innesii* exhibited the highest gene content, averaging 3,489 genes (range: 3,064–4,069), followed by *E. entomosocium* with 3,455 genes (3,086–3,810), and *E. casseliflavus* with 3,370 genes (2,916– 3,782). The Kruskal-Wallis test evidenced the existence of at least one pair of groups with a statistical difference in the median of the numbers of genes (p-value: 0.0373). The post-hoc test indicates a significant difference between the median number of genes of *E. casseliflavus* and *E. innesii* (adjusted p-value: 0.0477).

As representants of phylum Firmicutes, enterococci are Gram-positive bacteria with low GC content (ranging from 34% to 45%) (García-Solache and Rice 2019; Schwartzman et al. 2024). Compared to that of ancestrally related *Vagococcus* (33%), all four enterococci clades exhibit an increase in mean G+C content. *Enterococcus* Clade I averaged 36.2% (+/−0.9%), Clade II averages 37.3% (+/−1.3%), Clade III averages 39.6% (+/− 1.4%), and Clade IV averages 39.8% (+/− 2.7%) GC (Schwartzman et al. 2024). In this study, as expected to Clade IV species, *E. casseliflavus* exhibited the highest average GC content (42.44%; range: 40.3–43.31%), followed by *E. innesii* (42.16%; 40.86–42.74%) and *E. entomosocium* (41.89%; 40.26–42.83%) (Figure 3C). A total of seven samples (two from *E. casseliflavus*, three from *E. entomosocium*, and two from *E. innesii*) were considered to be outliers in the box plots, all of them showing GC content below 41.8%. The Kruskal-Wallis test evidenced the existence of at least one pair of groups with different median GC content (p-value: 1.61e-05). Post-hoc Dunn’s test indicates that the GC content median of *E. casseliflavus* differs significantly from both *E. entomosocium* (adjusted p-value: 0.00498) and *E. innesii* (adjusted p-value: 0.0002).

Pangenome analyses indicated that all three species possess open pangenomes, suggesting that the number of genes continues to expand with the addition of new genomes (Figures 3E, F, and G). This pattern has been widely reported in other *Enterococcus* species, including *E. faecium* (Lebreton et al., 2013) and *E. raffinosus* (Sharon et al., 2023). The *E. casseliflavus* pangenome (Figure 3E) comprised 8,614 genes, including 2,060 core genes, 4,940 accessory genes, and 1,650 strain-specific genes. Pangenome expansion followed the parameters *k* = 3188.39 and γ = 0.21, with a strong model fit (mean R² = 0.99; 95% CI = [0.95, 1.00]). Core genome contraction was described by *c* = 3012.16 and δ = −0.08 (mean R² = 0.87; 95% CI = [0.61, 0.96]). The *E. entomosocium* pangenome (Figure 3F) consisted of 5,167 genes, distributed into 2,627 core genes, 1,597 accessory genes, and 943 strain-specific genes. The pangenome growth parameters were *k* = 3287.65 and γ = 0.17 (mean R² = 0.93; 95% CI = [0.81, 0.98]). Core genome reduction was modeled with *c* = 3211.29 and δ = −0.08 (mean R² = 0.77; 95% CI = [0.01, 0.96]). The *E. innesii* pangenome (Figure 3G) encompassed 6,677 genes, including 2,398 core genes, 3,158 accessory genes, and 1,121 strain-specific genes. Pangenome expansion was characterized by *k* = 3442.87 and γ = 0.20 (mean R² = 0.95; 95% CI = [0.87, 0.99]), whereas core genome reduction followed *c* = 3059.37 and δ = −0.08 (mean R² = 0.79; 95% CI = [0.64, 0.92]). The open nature of these pangenomes is largely driven by a high proportion of accessory genes, which can be frequently exchanged through horizontal gene transfer, thereby facilitating rapid adaptation to diverse environments (McInerney et al. 2017). This genomic flexibility likely reflects the ecological versatility of these species, which are known to inhabit a broad range of niches.

### Genes over- and under-represented among species

We identified genes with species-specific patterns of presence or absence (Table S6) by analyzing the gene presence/absence matrix generated by Orthofinder (Table S5). A gene was considered *over-represented* in a species if it was present in at least 85% of its isolates and absent from at least 85% of isolates from each other species. Conversely, a gene was considered *under-represented* in a species if it was absent from at least 85% of its isolates but present in at least 85% of isolates from each remaining species. Orthogroups identified using these criteria were subsequently manually curated to determine whether multiple orthogroups corresponded to the same biological gene family. This assessment was based on synteny, functional annotation, and sequence similarity. As result, we detected nine, seven, and one over-represented genes in *E. entomosocium*, *E. casseliflavus*, and *E. innesii*, respectively. In *E. casseliflavus*, over-represented genes included *BglA*, *BglF*, and *BglG*, which are associated with the uptake and metabolism of β-glucosides. Among the genes over-represented in *E. entomosocium*, five form a putative operon encoding enzymes involved in ribose metabolism, such as ADP-ribosyltransferase and ADP-ribose hydrolase. It is noteworthy that *E. casseliflavus* and *E. flavescens* were originally described as separate species based on their ability to ferment ribose, which is present in the former but absent in the latter (Pompei et al, 1992). However, a closer examination of these data reveals that the genes associated with ribose fermentation in *E. casseliflavus* are restricted to the type strain (DSM 20680) and a small group of closely related isolates (FDAARGOS 1120, JCM 8723, NBRC 100478, and NCTC 12361). This finding suggests that *E. casseliflavus* does not typically possess the ability to ferment ribose, and that this trait should instead be considered an exception rather than a defining characteristic of the species. Regarding under-represented genes, we identified none in *E. entomosocium*, three in *E. casseliflavus*, and eight in *E. innesii*. Notably, five genes under-represented in *E. innesii* (*fabD*, *madA*, *mdcC*, *accD*, and *mdcG*) form an operon involved in malonate metabolism.

The presence of genes associated with carbohydrate metabolism suggests the ability to utilize these compounds as carbon sources, potentially conferring ecological advantages in specific niches and hosts, a trait commonly observed in *Enterococcus* species (Lebreton et al. 2013; Lebreton et al., 2017; Schwartzman et al. 2024). *E. cecorum* and *E. columbae*, for example, two closely related species also from clade IV, exhibit distinct ecological distributions, with *E. cecorum* occurring across a wide range of avian hosts (Aarestrup et al., 2002), whereas *E. columbae* appears to be largely restricted to pigeons (Devriese et al., 1990; Baele et al., 2002). Comparative genomic analyses suggested that *E. columbae* has diverveged from a common ancestral species with *E. cecorum* and acquired 35 genes involved in carbohydrate metabolism, enabling the utilization of plant-derived substrates such as pectin and rhamnogalacturonan I (RG-I), which likely supports its adaptation to the plant-rich diet of Columbidae birds and contributes to its ecological specialization and divergence (Podulka, S. 2004; Lebreton et al., 2017).

### Pathogenic potential of E. casseliflavus and related species

We identified 14 distinct genes associated with resistance to biocides and tolerance to metals (Figure 4A, Table S7). On average, isolates carried 2.16 genes from this category, with *E. casseliflavus*, *E. innesii*, and *E. entomosocium*, presenting mean counts of 2.34, 1.86, 1.28 genes, respectively. The gene *chtR* was the most widespread, being present in nearly all isolates, except for a single *E. casseliflavus* isolate (V387). A total of 97 isolates carried *only* the *chtr* gene, including 65 from *E. casseliflavus*, 19 from *E. innesii*, and 13 from *E. entomosocium*. The *chtR* gene has been described as a component of the *chtRS* system in *E. faecium*, where it contributes to chlorhexidine tolerance. However, in contrast to *E. faecium*, the three species analyzed here lack the *chtS* gene, which is reported to be required to confer the tolerance (Prieto et al., 2017). The second most prevalent group comprised the *tcrABYZ* genes, associated with copper resistance. Isolates were largely divided into those that carried at least one gene (42 *E. casseliflavus*, one *E. entomosocium*, and six *E. innesii*) and those that did not. Notably, the *E. casseliflavus* isolate FDAARGOS_1122 harbored the highest number of resistance genes in this category (12), including *chtr*, *tcrABYZ*, and the operon containing genes involved in mercury tolerance (*merA*, *merB1*, *merB2*, *merB3*, *merP*, *merR1*, *merR2*). Additional resistance genes included *arsB* (detected in 8 *E. casseliflavus* and 5 *E. innesii* isolates) and *cadC* (found in 4 *E. casseliflavus* isolates).

**Figure 4:**
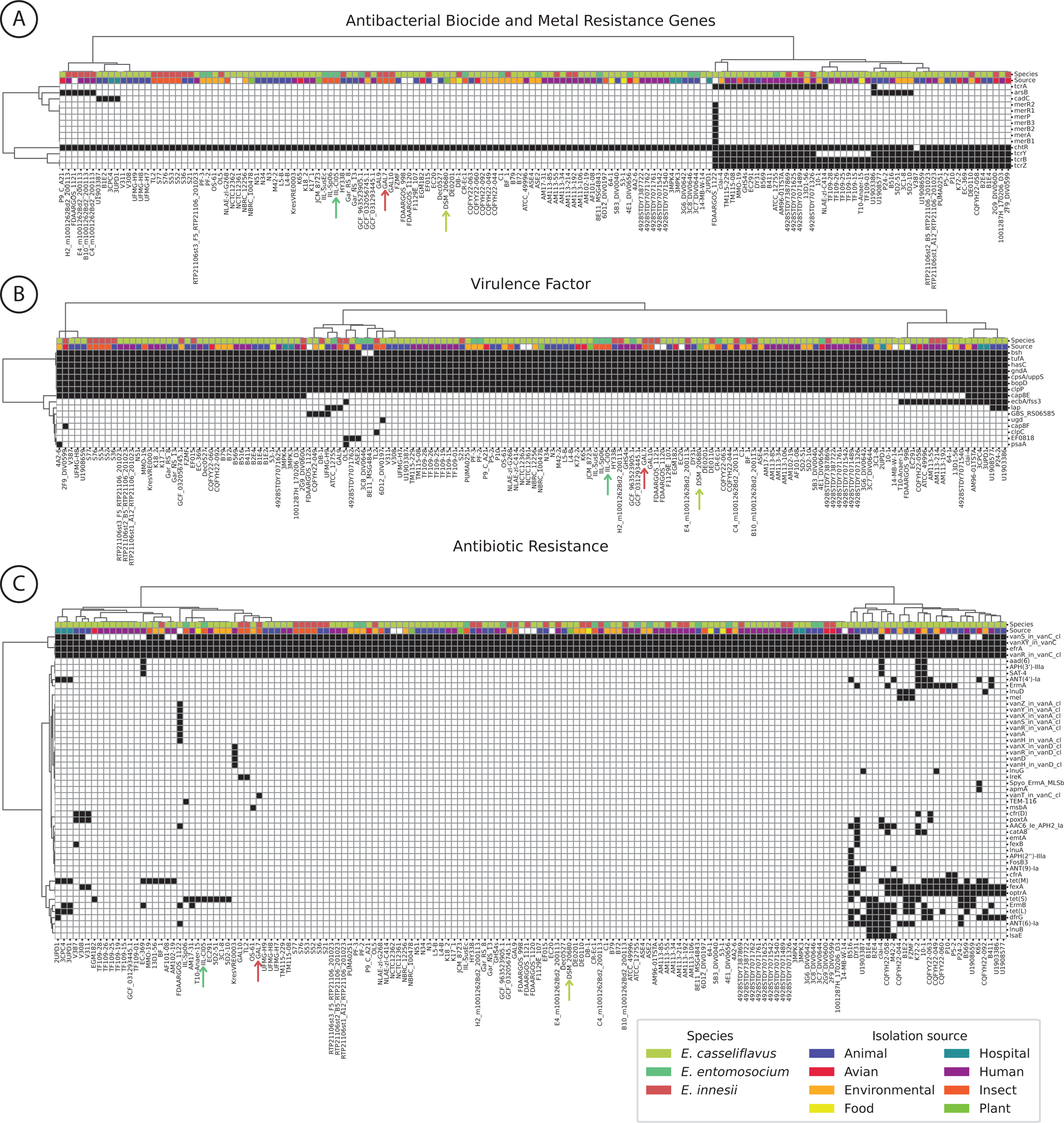
Distribution of virulence and resistance genes among the three *Enterococcus* species. Hierarchical clustering and presence/absence matrices of **(A)** antibacterial biocide and metal tolerance-associated genes, **(B)** virulence, and **(C)** antibiotic resistance genes. Columns and lines represent samples and genes, respectively. Filled cells in the matrix denote the presence of the gene. The first two lines of the matrices indicate the species and the isolation source of each sample. Arrows below the matrices indicate the positions of the reference strains of the species *E. casseliflavus* (DSM 20680, yellow), *E. entomosocium* (IIL-Cl05, green), and *E. innesii* (GAL7, red). More details about the genes are available in Table S7.

A total of 16 genes associated with potential virulence traits (Figure 4B, Table S7) were also identified, with isolates carrying an average of 7.53 of such genes. The mean number of virulence genes was similar among species: 7.63 in *E. innesii,* 7.57 in *E. casseliflavus*, and 7.0 in *E. entomosocium*. Seven of these genes (*bsh*, *tufA*, *hasC*, *gndA*, *cpsA/uppS*, *bopD*, *clpP*) were found in all or nearly all isolates. Notably, several of them (*bopD, bsh, cpsA/uppS,* and *hasC*) are also commonly found in *E. faecium* (Sanderson et al., 2022), suggesting that they may represent broadly conserved functions within the genus. The remaining virulence genes were exclusive to *E. casseliflavus* and *E. innesii*. The gene *cap8E* was present in 48 isolates, including 39 *E. casseliflavus* and 9 *E. innesii*. The gene *ecbA/fss3* was identified in 15 *E. casseliflavus* and 4 *E. innesii* isolates.

Finally, we detected 50 distinct antibiotic resistance genes across all isolates (Figure 4C, Table S7), with a mean of 5.45 genes per genome. On average, *E. casseliflavus*, *E. entomosocium*, and *E. innesii* harbored 5.96, 4.21, and 4.13 resistance genes, respectively. *E. casseliflavus* stood out by carrying the largest diversity of resistance genes (47 unique genes), compared to only 7 and 6 unique genes in *E. innesii and E. entomosocium* respectively. The isolate with the highest number of resistance genes was B516 (*E. casseliflavus*), which harbored 17 genes. All genomes carried *vanXY*, *vanR* (components of the *vanC* cluster), and *efrA*, while *vanS* was present in the majority (131/156) of strains. Two strains showed unique resistance profiles. FDAARGOS_1122 (*E. casseliflavus*) was the only strain harboring genes from the *vanA* cluster (*vanAHRSXYZ*), which confer high-level resistance to vancomycin and is commonly found in *E. faecium*. In contrast, KresVRE0003 (*E. casseliflavus*) was the only isolate with the *vanD* cluster (*vanDHRX*), a relatively rare genotype that has been sporadically reported in *E. faecium*, *E. faecalis*, *E. gallinarum*, *E. avium*, and *E. raffinosus* (Depardieu et al., 2009), and more recently in *E. casseliflavus* (Rubaye et al., 2021). Additionally, a subset of 26 *E. casseliflavus* isolates formed a distinct cluster characterized by a high number of antibiotic resistance genes. Among the most frequently detected resistance genes were *tet(M)* (24 isolates), *optrA* (22), *fexA* (22), *tet(S)* (19), *tet(L)* (17), and *ermB* (17), suggesting the widespread presence of resistance mechanisms against tetracyclines and macrolides.

### Species with host-associated ecological niches

We retrieved metadata on the source of isolation for each sample from the NCBI database to assess potential species-specific patterns. Of the 156 genomes, only seven lacked this information. The remaining 149 genomes were classified into eight categories: mammal, avian, environmental, food, hospital, human, insect, and plant. Five of these categories were represented across all three species. The food category was absent in *E. entomosocium*, whereas hospital and plant sources were observed only in *E. casseliflavus*. The absence of certain categories in *E. entomosocium* and *E. innesii* likely reflects their smaller sample sizes, suggesting no clear species-specific association with isolation source. We also verified that samples from the same source of isolation do not cluster into monophyletic groups in the inferred phylogenetic tree (Figure 2), indicating that the trait in question likely arose multiple times independently during the evolution.

We also conducted an association analysis between the isolation source and some genomic features, including the number of genes (Figure 5A), genome size (Figure 5B), and GC content (Figure 5C). In terms of gene number, all groups showed an average of 3,304-3,407 genes, except for samples isolated from insects, which exhibited a higher average of 3,721 genes. Similarly, the average genome size ranged from 3.49 to 3.61 Mb across groups, with the insect group again standing out, with a larger average genome size of 3.84 Mb. Regarding GC content, the lowest average was observed in the avian group (41.95%), followed closely by the insect group (42.07%), while the other groups displayed averages ranging from 42.28% to 42.48%.

**Figure 5:**
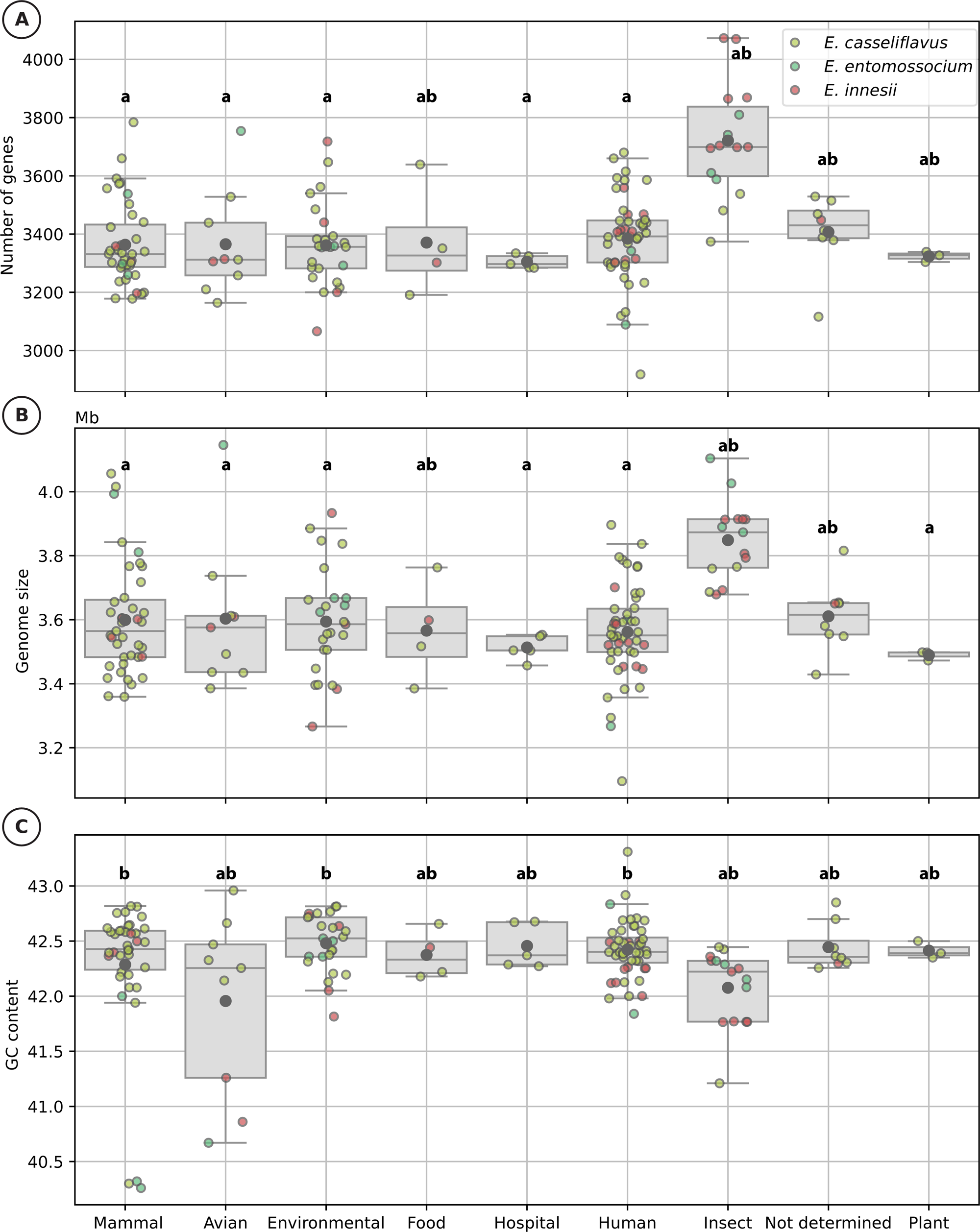
Genomic feature comparisons across isolation sources. **(A–C)** Boxplots showing the distribution of **(A)** number of genes, **(B)** genome size (Mb), and **(C)** GC content (%) across different isolation sources. Large gray circles represent the mean values for each group. Letters above the boxes indicate statistically significant groupings based on Dunn’s test with Holm correction (q-value < 0.05). Data points are colored according to species: *E. casseliflavus* (yellow), *E. entomosocium* (green), and *E. innesii* (red).

Statistical analysis using the Kruskal-Wallis test revealed significant differences among group medians for all genomic features analyzed. Post hoc comparisons with the Dunn test further indicated that all significant differences involved samples isolated from insects. Specifically, insect-derived samples had significantly higher numbers of genes and larger genome sizes compared to other groups, while their GC content was significantly lower. This pattern may reflect the acquisition of accessory genetic material with adaptation to insect-associated ecological niches, potentially increasing metabolic versatility and environmental fitness.

## Conclusions

Vancomycin-resistant strains of *E. faecalis* and *E. faecium* emerged as major causes of multidrug-resistant hospital-associated infections in the mid-1980s, with resistance typically acquired through mobile genetic elements. The success of these species as nosocomial pathogens highlights the critical role of gene exchange in the evolution and dissemination of antimicrobial resistance within the genus. In contrast, other enterococcal species, including *E. casseliflavus* and closely related species, as we showed here, harbored the *vanC* operon, which is associated with low-level vancomycin resistance. Although they are less frequently associated with human infections, their broad ecological distribution and genomic plasticity suggest that they may serve as overlooked reservoirs of resistance and mobile genetic elements that could be mobilized into the clinical environment. Furthermore, we provide a comprehensive genomic reassessment of *Enterococcus casseliflavus* and closely related taxa, revealing substantial inconsistencies in current species designations in public databases. By integrating multiple genome-based approaches, we demonstrate that these methods are essential for accurate species delineation within this group. The strong concordance across analyses consistently resolves these enterococci into three well-defined clusters using a higher discrimination threshold (ANI >96.6): one comprising *E. casseliflavus* and *E. flavescens*, supporting their synonymy, and two additional clusters corresponding to *E. innesii* and *E. entomosocium*, which are clearly delineated as distinct species. Our results also showed that these lineages harbor extensive genetic diversity and a rich repertoire of accessory genes, including putative resistance determinants, underscoring their potential role as reservoirs of antimicrobial resistance. Given the capacity of enterococci to exchange mobile elements, clarifying their genetic diversity is a critical step toward improving surveillance and mitigating the spread of antimicrobial resistance.

## CRediT authorship contribution statement

**Matheus Miguel Soares de Medeiros Lima:** Conceptualization, Methodology, Formal analysis, Writing – original draft. **Janira Prichula:** Conceptualization, Methodology, Data curation, Writing – original draft, Writing – review & editing. **Tetsu Sakamoto:** Data curation, Writing – review & editing, Project administration, Funding acquisition.

## Acknowledgment

We thank the teams from Bioinformatics Multidisciplinary Environment (BioME/IMD) at UFRN, and High-Performance Computing Center (NPAD) at UFRN for the support on the computational resource. We are also grateful to NCBI teams for the curation and maintenance of the data used in this work.

