## Supplementary Fig. 1 for "Resolving Taxonomic Boundaries and Revealing Genetic Diversity in *Enterococcus casseliflavus* and Closely Related Taxa"

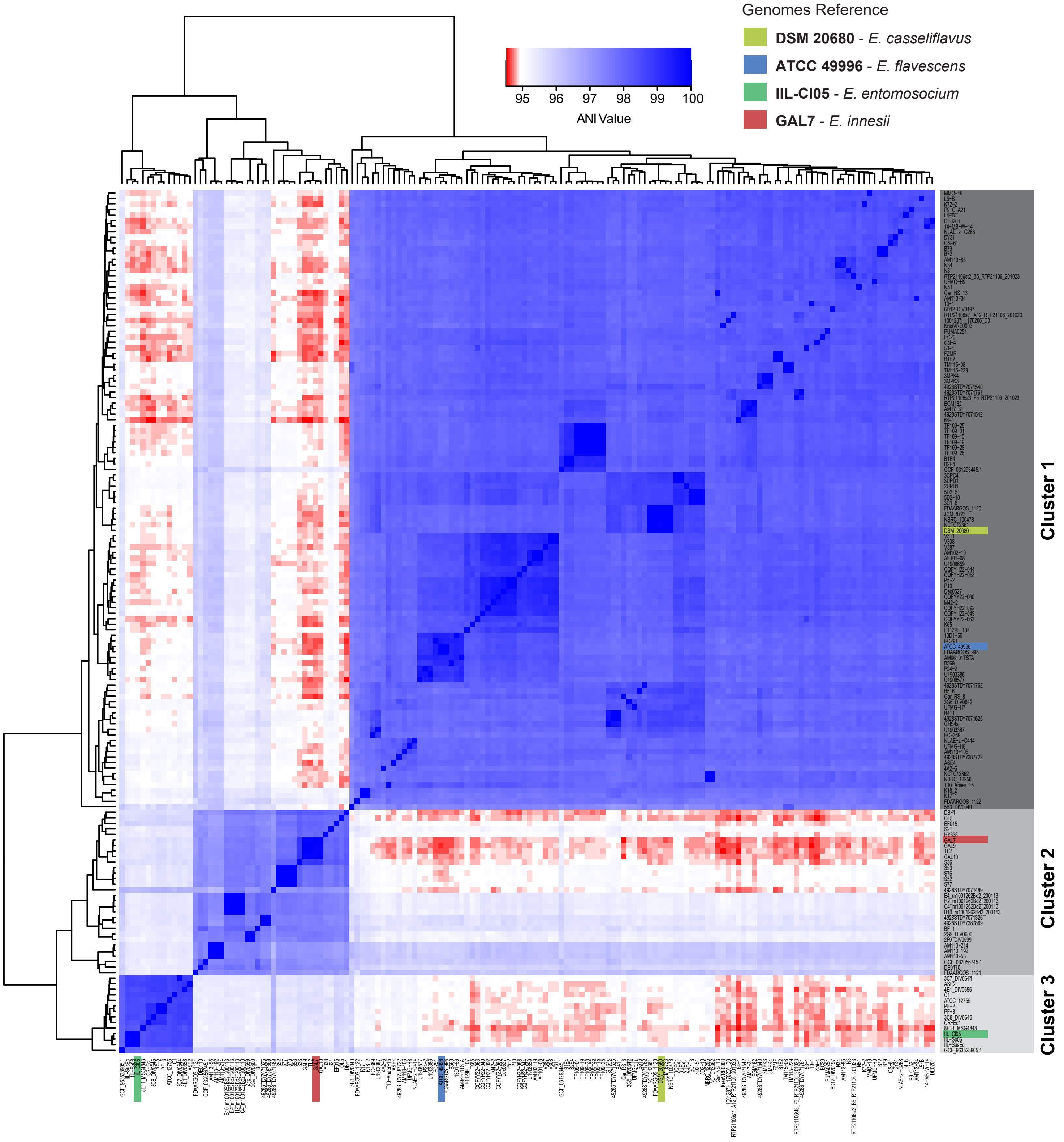


**Supplementary Fig. 1: Heatmap displaying the ANI values of the 156 selected *Enterococcus* genomes**, generated using FastANI. The figure illustrates the delineation of three distinct clusters. Type strain genomes are highlighted in blue (*E. flavescens* ATCC 49996), red (*E. innesii* GAL7), yellow (*E. casseliflavus* DSM 20680), and green (*E. entomosocium* IIL-Cl05) stars. Data used to generate this figure can be found in Table S2.
